# Neuromodulation enhances the capability and efficiency of spiking neural networks

**DOI:** 10.1101/2025.07.25.666748

**Authors:** AbdelQader AlKilany, Dan F. M. Goodman

## Abstract

Spiking neurons underlie the brain’s extreme energy efficiency, and therefore have great potential in neuromorphic computing, although realising this efficiency in practice has proven challenging. We use neuromodulation, a biological mechanism that lets the network dynamically and contextually modify its own parameters. We find it substantially increases performance across a range of sensory processing tasks, including a challenging new speech-in-noise dataset we introduce, with very few additional resources (neurons, energy, parameters). Neuromodulatory networks are space and energy efficient thanks to two mechanisms that are directly relevant to neuromorphic computing and biology: firstly, they allow for an order-of-magnitude reduction in the number of neurons required; and secondly, they enable very sparse firing, achieving better results using orders of magnitude fewer spikes. Together, these properties may throw light on the computational role of neuromodulation in biology, and make neuromodulation an ideal mechanism to improve performance and efficiency for neuromorphic devices.

---

It is widely believed that the discrete spiking mechanism of biological brains is critical for overall energy efficiency thanks to its noise robustness in long-range communication [Laughlin et al., 1998, Attwell and Laughlin, 2001, Lennie, 2003]. The potential for energy efficient computation makes spiking neural networks very attractive for the design of neuromorphic computing devices and algorithms [Schuman et al., 2022]. However, in practice there is a practical limit to the scaling of these systems, both algorithmically [Neftci et al., 2019, Bellec et al., 2020, Eshraghian et al., 2023], and in terms of the number of neurons, synapses and events that can be processed [Kudithipudi et al., 2025, Davies et al., 2021]. Although ultimately these fundamental issues in learning and scaling remain to be solved, we can make progress towards practical neuromorphic systems by improving their performance within a fixed budget (of computation, area, power, etc.).

One approach to improving the capability of spiking neural networks within a fixed budget is to draw inspiration from biological mechanisms that increase capacity without significantly increasing computational complexity. Examples include heterogeneity [Perez-Nieves et al., 2021, Dahmen et al., 2026], adaptation [Bellec et al., 2018, Salaj et al., 2021], delays [Sun et al., 2023, 2025] or dual memory pathways [Sun et al., 2026]. Here, we extend these investigations to include the role of neuromodulation, which we will see can act as a sort of attentional mechanism, which has been shown to be very powerful in machine learning [Bahdanau et al., 2014, Vaswani et al., 2017].

In addition to fast neurotransmitters acting at synapses, some neurons release neuromodulators that can change properties of groups of cells [Nadim and Bucher, 2014, Mei et al., 2022]. These are usually studied at slow timescales, for example looking at their possible computational roles in learning [Doya, 2002, Grossman and Cohen, 2022]. Recent evidence, however, has suggested they may be important in sensory processing at faster time scales, even sub-second [Hangya et al., 2015, Bang et al., 2020]. A complete picture of their role remains elusive, particularly at fast timescales, but the evidence is suggestive enough to warrant investigation of its potential computational role in these settings. Simplified models of neuromodulation in spiking neural networks have been suggested before in a neuromorphic context [Krichmar, 2012, Ribar and Sepulchre, 2019] and in differentiable models of plasticity [Schmidgall et al., 2021]. However, a detailed study of their potential computational roles in augmenting the computational capabilities of spiking neural networks is lacking.

Here, we investigate how an abstract, general model of neuromodulation can augment spiking neural networks. In brief, we find that it enhances sensory processing across a range of challenging tasks, and has a particularly salient role in handling temporally complex signals. These mechanisms appear to be robust and general, and are therefore likely to be valuable in understanding the role of neuromodulation in the brain, and in the design of efficient neuromorphic devices.

## 1 Results

We started from the hypothesis that allowing networks of spiking neurons to dynamically adjust their own parameters in a context-sensitive way would improve their performance on tasks, particularly those with rich temporal structure (as in Perez-Nieves et al. 2021). To test this, we started from a simple baseline spiking neural network (SNN) consisting of a layer of spiking input neurons connected to a recurrent hidden layer of spiking neurons, which was in turn connected to a linear readout layer. The hidden layer of spiking neurons includes trainable parameters such as time constants, resting potentials and thresholds. In addition, all synaptic weights between layers were trainable. For training, we used the method of surrogate gradient descent [Neftci et al., 2019]. We used three datasets, including two spiking speech recognition datasets (SHD and SSC; Cramer et al. 2022), and one visual gesture recognition dataset based on a neuromorphic event camera (DVS128 Gestures; Amir et al. 2017). For our baseline model, we use 256 hidden neurons, as larger networks provide only marginal performance improvements while requiring substantially more computation and training time.

### 1.1 Neuromodulation enhances sensory processing

In our first model of neuromodulation, we add an artificial neural network that we refer to as the *neuromodulatory network*. It accumulates inputs from the hidden layer of the spiking network for a fixed duration (*K* timesteps), and passes this input through a multilayer perceptron (MLP). The outputs of this MLP are used as the new parameter values (time constants, threshold, etc.) for the next simulation window of *K* timesteps. The whole process then repeats for the remainder of the input stimulus (fig. 1A,D). We find that this leads to large improvements in performance across all datasets (fig. 1C,F,I). SHD and SSC benefited most from short update intervals, whereas DVS peaked at a longer interval, consistent with a trade-off between rapid parameter adaptation and integrating enough recent context to make a useful modulatory update.

**Figure 1:**
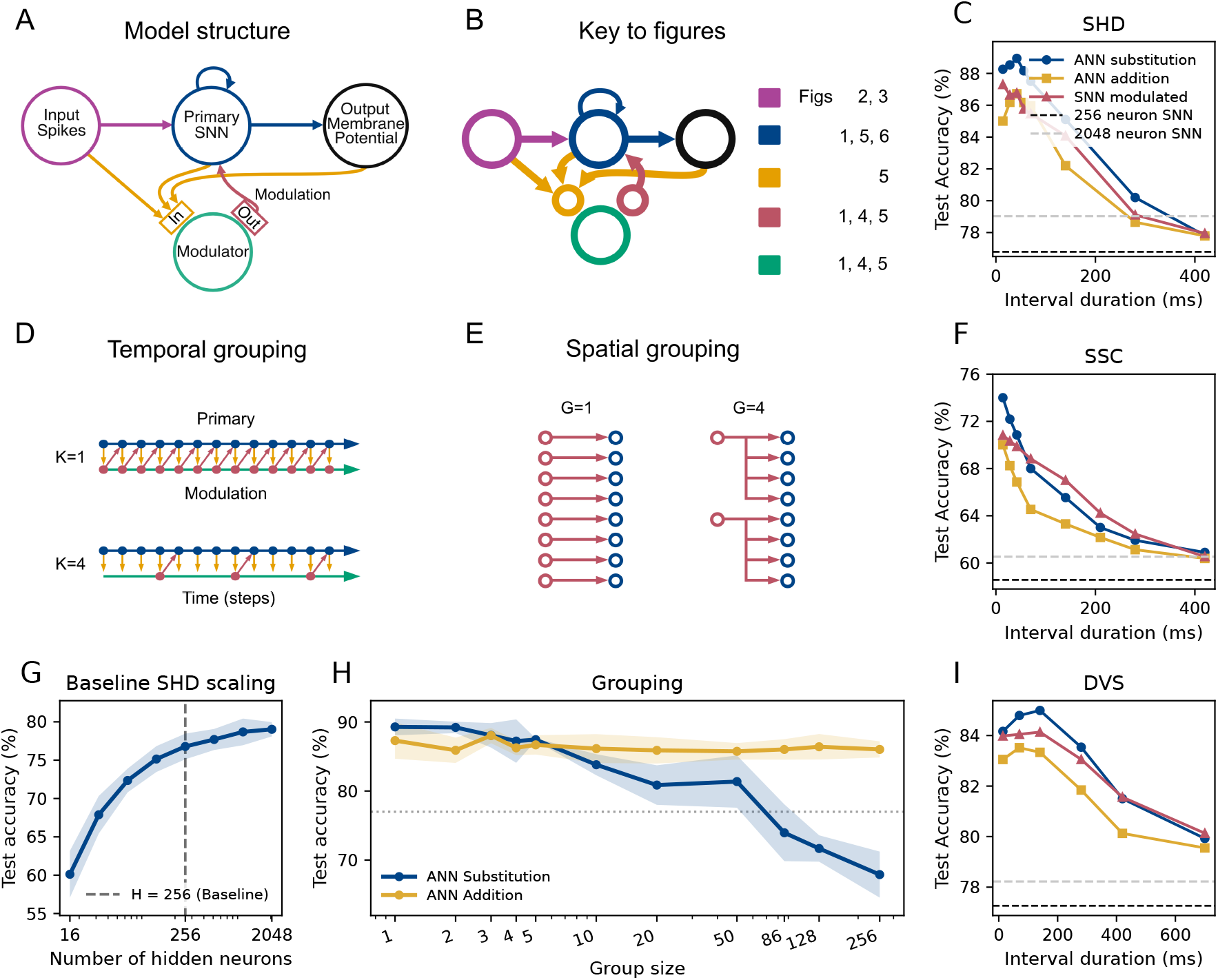
Neuromodulation enhances sensory processing in spiking neural networks. **(A)** Model structure. The primary spiking neural network (SNN) processes input spike trains, while a secondary modulator network adjusts parameters of the primary network based on the context. **(B)** Key to subsequent figures. **(C,F,I)** Test accuracy on SHD, SSC and DVS128 Gestures, respectively, as a function of the update interval of the modulator network. Results are shown for ANN substitution and addition methods, and with an SNN as modulator. Dashed horizontal lines indicate the mean accuracies of unmodulated networks. **(D)** Temporal grouping, showing when the modulator network receives inputs from and modifies the primary network at two different modulation intervals. **(E)** Spatial grouping, showing that a single neuromodulator output can affect a single neuron or group at different grouping sizes. **(G)** Scaling performance for unmodulated SNNs with the number of hidden neurons on the SHD task. The dashed vertical line marks the 256-hidden-neuron baseline used in most experiments. Shaded regions indicate mean *±* one standard deviation across runs. **(H)** Effect of spatial grouping size on performance for neuromodulated SNNs on the SHD task. The dotted line shows the performance of the unmodulated SNN, and shaded regions indicate variability across runs.

To separate the effect of neuromodulation from the effect of simply scaling the primary SNN, we also trained unmodulated SNNs with increasing hidden-layer sizes. Direct scaling improved SHD performance, but did not reproduce the gains achieved by modulation over the tested parameter range (fig. 1G). This indicates that the main performance improvements are not explained solely by adding more trainable parameters.

Next, we tested performance when the neuromodulatory network can only modify (increase or decrease) existing parameters rather than replacing them. This may be more plausible as a biophysical model and potentially more compressible in a neuromorphic context. We find that in this case, performance is still improved in all conditions, but not as much as when parameters are replaced (fig. 1C,F,I).

The effects of neuromodulation in biology can be felt across a wide range of spatial scales. In our model, these effects can be different for each single modulated neuron, or they may be shared across groups of neurons. Performance of the addition-based modulation remained comparatively stable at all grouping sizes. However, substitution-based modulation only performed well for small and intermediate groups, degrading below the performance of the unmodulated reference when a single modulatory value was broadcast to many neurons (fig. 1H). This happens because at large group sizes substitution forces the network to be homogeneous (whereas the unmodulated network is heterogeneous).

### 1.2 Temporally dynamic gain control improves effective signal to noise ratio

Neuromodulation allows the network to change its computational properties dynamically in time within a stimulus, akin to dynamic gain control. To test this capability in a more challenging setting, we created a speech-in-noise variant of SHD [AlKilany and Goodman, 2026] by adding either sinusoidally amplitude-modulated (SAM) noise or recordings of natural background noise. Neuromodulation substantially improved robustness to both SAM and natural noise (fig. 2A,B). The largest gains in accuracy (15-18% higher) occurred around the transition between chance performance and reliable recognition (-17 dB SNR in SAM noise, -5 dB SNR in natural noise).

**Figure 2:**
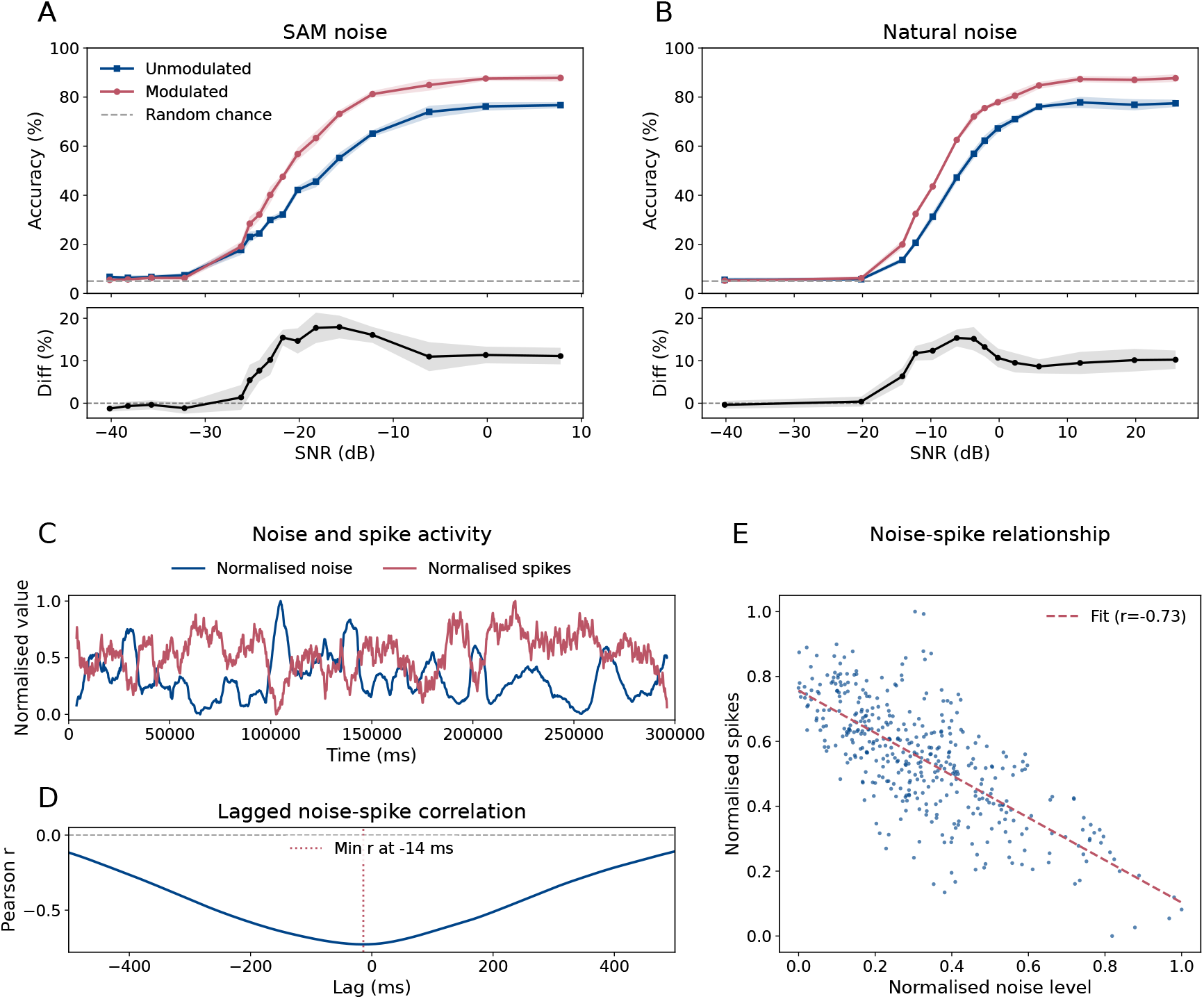
Neuromodulation improves robustness to speech in noise. **(A,B)** Robustness to sinusoidally amplitude-modulated (SAM) noise and natural cafe noise across signal-to-noise ratios (SNRs). Top panels show classification accuracy for unmodulated and modulated networks, with chance performance indicated by the dashed line. Bottom panels show the absolute accuracy difference between the modulated and unmodulated networks. **(C)** Smoothed and normalised background noise level and average spike activity over time. Network activity increases during quieter periods, indicating an inverse relationship between spike output and natural noise level. **(D)** Lagged Pearson correlation between spike activity and noise level, showing that the strongest modulation occurs with a delay of approximately 14 ms. **(E)** Normalised spike activity plotted against noise level for sampled time windows, showing a strong negative correlation between noise level and network output.

It has been speculated that the human auditory system is able to “listen in the dips”, focusing processing on brief moments where the noise is low relative to the speech signal [Peters et al., 1998, Lorenzi et al., 2006]. This is an ability that even state-of-the-art machine listening systems still struggle with [O’Shaughnessy, 2024]. Since the modulated network has the capability to do dynamic gain control (by increasing the firing threshold for example), we tested whether it implements a dip-listening strategy. Our networks fire at a higher rate when the noise level is higher (fig. 3B,E) because the background noise leads to many more input spikes. We therefore computed the ratio of spike rates in the hidden layer to spike rates in the input, and find that this ratio is higher at dips in the noise as would be expected if the network is doing dip-listening (fig. 3C). Both modulated and unmodulated networks show a negative relationship between the noise level and the hidden/input firing-rate ratio, however the relationship is substantially stronger in the modulated network (fig. 3H). Furthermore, by inspecting the distribution of modulated parameters, we can see changes consistent with dip-listening, e.g. a large dip in the post-spike reset value when the noise level is high (making it harder for the neuron to fire multiple spikes; fig. 3D).

**Figure 3:**
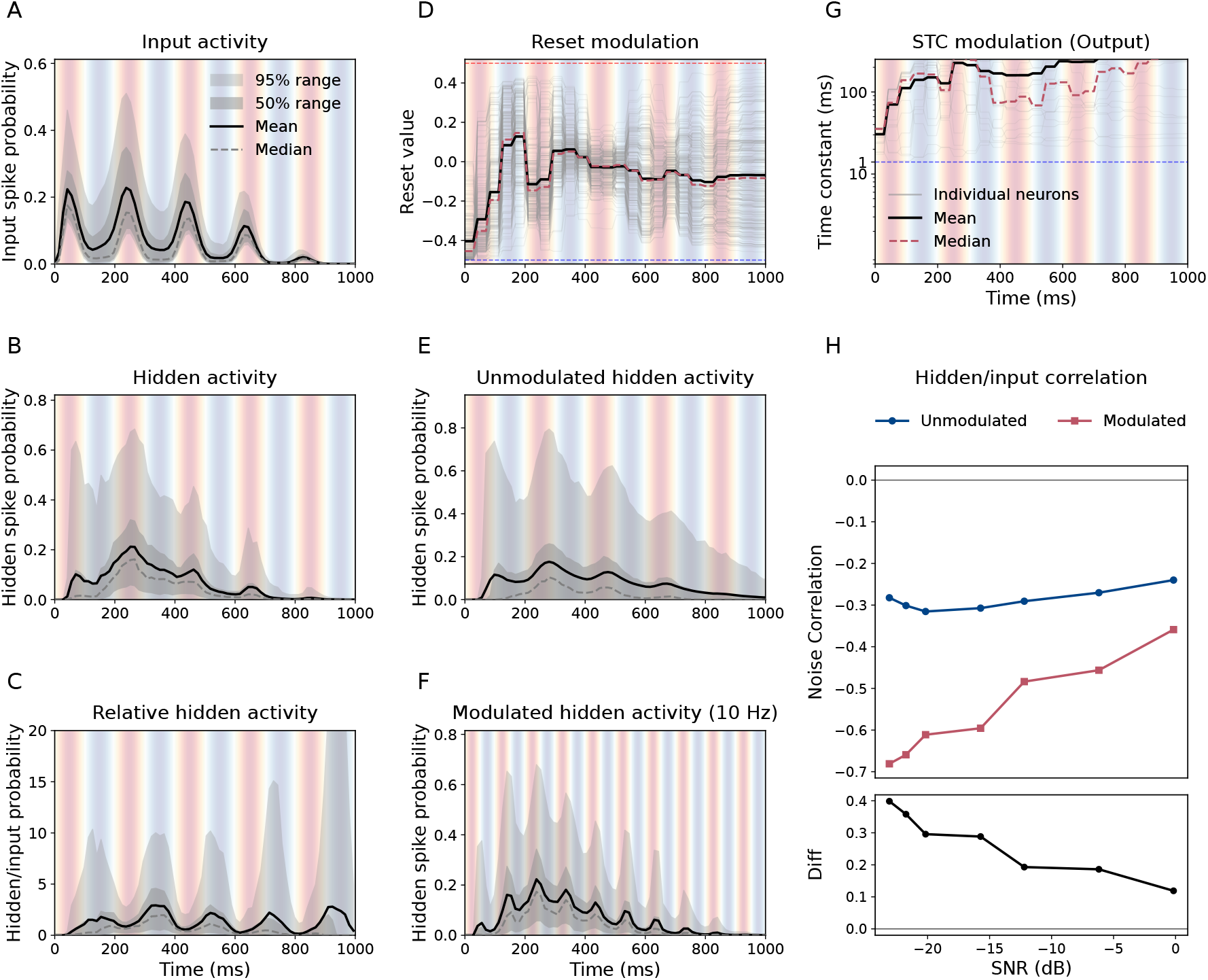
Time-resolved activity and parameter modulation under sinusoidally amplitude-modulated noise. **(A)** Input spike activity for SHD with 5 Hz sinusoidally amplitude-modulated background noise at -14 dB SNR; blue indicates low noise amplitude and red indicates high noise amplitude. **(B)** Hidden activity of the modulated network in the same condition. **(C)** Relative hidden activity, computed as hidden activity divided by input activity. **(D)** Reset parameter modulation in the modulated network. **(E)** Hidden activity of the unmodulated network under the same noise condition. **(F)** Hidden activity of the same modulated network trained at 5 Hz and tested on 10 Hz modulation at the same noise amplitude. **(G)** Output-layer beta modulation in the modulated network. **(H)** Correlation between hidden to input firing rate and noise amplitude across SNR for 10 Hz sinusoidally modulated noise. Top: noise correlation for unmodulated and modulated networks. Bottom: gap between unmodulated and modulated networks, computed as unmodulated minus modulated correlation.

To test whether this was an adaptive strategy rather than memorisation of a specific noise waveform, we randomised the initial phase of the SAM noise (the plots show a fixed initial phase for illustrative purposes). Further, we tested frequency generalisation by training at one modulation frequency and testing at another (fig. 3F). Finally, we see the same qualitative effect in unpredictable natural background noise: network activity increased during quieter moments and showed a negative relationship with the noise envelope (fig. 2C,E). The correlation between noise and activity is most strongly negative at a lag of around -14 ms (fig. 2D), suggesting that the network is able to cope with rapid changes in noise level.

### 1.3 Biologically constrained neuromodulation can simplify models and reduce parameters

Neuromodulation is carried out by a class of neurons that release a quantity of one or more of a small set of neurotransmitters, that then diffuse slowly over a wide area. We tested whether the performance gains of neuromodulation persist under four additional constraints: (i) slow drifting changes in parameter values rather than instantaneous updates (mimicking the slow diffusion process), (ii) limiting the number of distinct “neuromodulator types” available (mimicking the different types of neurotransmitters), (iii) dispersing modulation effects across overlapping groups of neurons (mimicking the release sites), and (iv) replacing the artificial controller with a spiking controller (since neuromodulatory neurons are also spiking neurons). In addition to mimicking biology, these changes have the potential to simplify the implementation and reduce the number of parameters.

#### 1.3.1 Parameter diffusion

We model the slow diffusive change to parameters via an exponential drift towards a target set by the neuromodulatory network, with a diffusion factor *λ* corresponding to a diffusion time constant −d*t/* log(1 −*λ*). Smaller *λ* values correspond to slower dynamics, while *λ* = 1 corresponds to instantaneous update. Performance remained well above the unmodulated baseline across the full range of *λ* (fig. 4A,B). Accuracy was highest at intermediate factors, showing that diffusion can be computationally beneficial.

**Figure 4:**
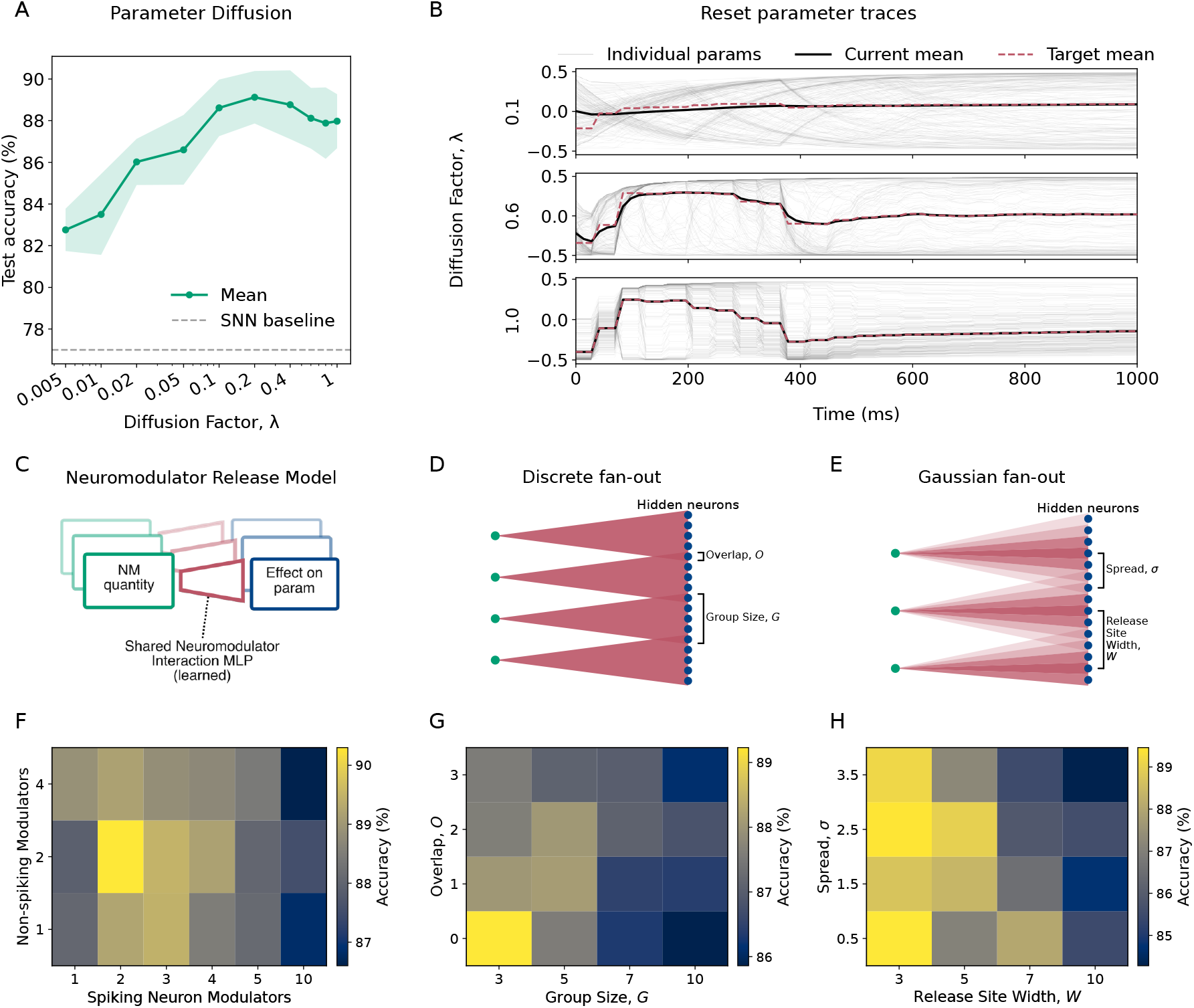
Biologically constrained neuromodulation preserves performance while simplifying control. **(A)** Test accuracy as a function of the diffusion factor applied to modulated parameters. Accuracy remains above the unmodulated SNN baseline across the full range and peaks at intermediate values. **(B)** Example reset-parameter trajectories under three diffusion factors (0.1, 0.6, 1.0; top to bottom). Smaller diffusion factors produce slower, more inertial parameter changes, whereas *λ* = 1.0 is instantaneous. **(C)** Schematic of the neuromodulator release model. The neuromodulatory network outputs a compressed set of neuromodulator quantities, which are then converted into parameter effects by a shared map. This enables coordinated parameter modulation while reducing the number of independent control signals required. **(D)** Discrete fan-out release architecture. Each neuromodulator release site targets a contiguous subset of hidden neurons. **(E)** Gaussian fan-out release architecture. Each neuromodulator release site produces a graded spatial profile across hidden neurons, with the strongest effects near the centre of the release site and weaker effects on more distant neurons. **(F)** Mean accuracy as a function of the number of neuromodulator types assigned to the spiking hidden population and the non-spiking readout population. Accuracy is shown as an image, with yellow indicating higher accuracy. **(G)** Accuracy under discrete spatial fan-out as a function of the width and overlap of the fan-out. **(H)** Accuracy under Gaussian spatial fan-out as a function of release-site width and Gaussian spread.

#### 1.3.2 Neuromodulator release model

Next, we constrained the neuromodulatory network to control target neuron parameters via a quantity of a small number of neuromodulator “types”. Since biological neuromodulators can have a range of effects on target neurons as well as interactions between neuromodulators, we model this by implementing a small trainable network with shared weights that converts the quantities of neuromodulators present into their effects on the parameters. Performance remained high with only a few neuromodulator types for both the spiking hidden layer and the non-spiking readout (fig. 4C,F), with accuracy only varying by around ±4%.

#### 1.3.3 Dispersion of neuromodulator effects

We then considered spatial dispersion where release sites target overlapping ranges of neurons with either a rectangular or Gaussian window. Both choices preserved high performance when release sites were relatively local, but accuracy declined for the widest and most overlapping effects (fig. 4E–H).

#### 1.3.4 Spiking modulator

So far, we have used an artificial neural network to control the parameters of a primary spiking neural network. We next tested the effect of replacing this artificial network with a spiking network, effectively making the network into a single SNN with different neuron types. In this case we cannot use the substitution method (since spiking neurons produce binary outputs not graded outputs), but something closer to the additive method: each spike of the modulator causes an increase or decrease in the value of some parameter. The spiking modulator reached broadly the same performance as the ANN modulator (fig. 1C,F,I).

### 1.4 Sparse modulation is an efficient strategy to reduce parameter counts

While neuromodulation improves performance, a naive neuromodulatory network can use many parameters, sometimes more than the number of parameters in the primary network. We therefore tested various strategies for reducing the size of the modulator while retaining most of the accuracy gains.

#### 1.4.1 Parameter reduction

##### Grouping

A simple way to reduce the size of the modulator network while retaining performance is to group multiple target neurons together (fig. 1H). This approximately divides the number of parameters by the grouping factor *G*.

##### Biological constraints

The biologically motivated constraints discussed above can also reduce modulator network size and parameter counts. Dispersion reduces the number of output channels, reducing parameters (fig. 4E–H). The neuromodulator release model takes this even further by grouping parameters (fig. 4C,D).

##### Input-channel compression

Following Cramer et al. [2022], we compressed the 700 SHD input channels using a fixed, non-learned aggregation. Adjacent channels were divided into non-overlapping groups and their spikes were summed independently at each time step. We compared three conditions (fig. 5A). In the unmodulated condition, the aggregated spike trains were supplied to the primary SNN. In the first modulated condition, the same aggregated spike trains were supplied to both the primary SNN and the modulator. In the second, compression was applied only to the modulator, while the primary SNN retained all the original channels. Compressing the primary SNN substantially reduced accuracy only at the strongest compression levels. By contrast, accuracy remained high across the range when only the modulator input was compressed, showing that the modulator required only a coarse summary of the sensory input while the primary SNN benefited from retaining its full frequency resolution.

**Figure 5:**
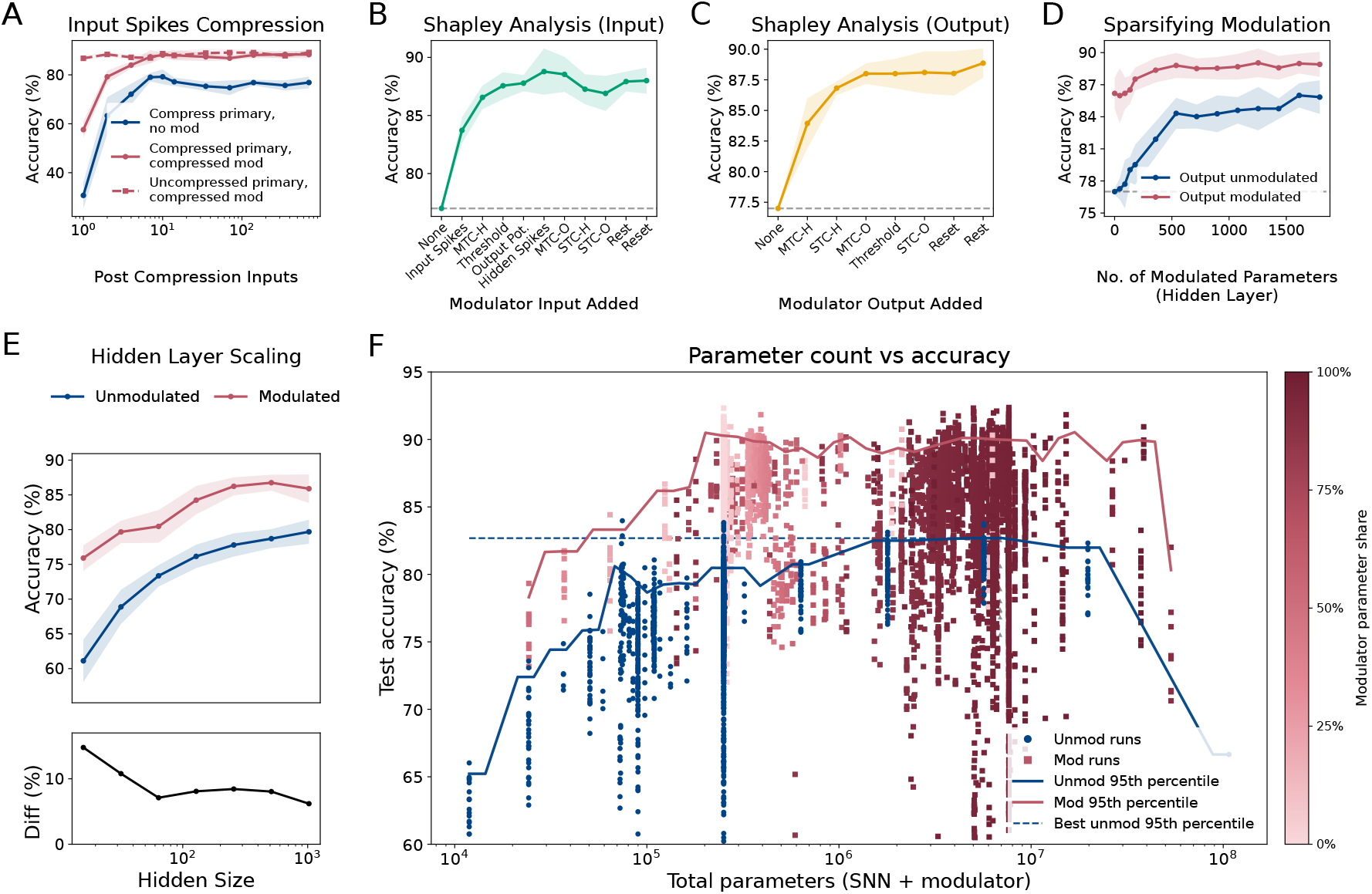
Sparse modulation reduces parameter counts while retaining performance. **(A)** Input compression. Blue shows an unmodulated SNN trained and tested with compressed primary inputs. Red solid shows a modulated network with the same compression applied to both the primary SNN pathway and the modulator. Red dashed shows modulator-only compression, where the primary SNN keeps full-resolution inputs. Left-to-right corresponds to decreasing compression. Shaded bands indicate mean *±* one standard deviation across runs. **(B,C)** Shapley analysis of modulator inputs and outputs. The left side of each figure indicates unmodulated performance when no inputs (B) or outputs (C) from the modulator network are present. Moving to the right, each step adds the next most important new input/output block so that at the right-hand side, all input/output blocks are present. The dashed line marks the unmodulated SNN baseline; shaded bands indicate mean *±* one standard deviation across runs. **(D)** Sparsifying modulation by limiting the number of hidden-layer parameters receiving modulation. Accuracy improves as more hidden parameters are modulated, and ensuring all outputs are modulated gives the strongest performance across the range. **(E)** Hidden-layer scaling for unmodulated and modulated SNNs. Top: accuracy versus hidden size. Bottom: modulation gain in percentage points, computed as modulated minus unmodulated accuracy. **(F)** Test accuracy as a function of total parameter count across all runs. Each point is a trained configuration, with unmodulated SNN runs shown separately from modulated runs. Solid curves show the 95th-percentile accuracy envelope over sliding windows in log-parameter space, summarising the best-performing configurations at each model scale. Point colour indicates the fraction of total parameters belonging to the modulator.

##### Disabling modulator pathways

In the base model, the modulator network receives spikes/potentials and current parameter values from the input, hidden and output layers, but not all of these are essential. We performed a Shapley analysis to rank which input features are most important (fig. 5B), and find that as long as the modulator network receives input spikes, hidden layer membrane time constants and thresholds, performance is more or less ideal. We did the same analysis on the outputs, excluding some parameters from neuromodulation (fig. 5C) and find that the key parameters are the hidden layer membrane and synaptic time constants, and the output layer membrane time constant.

##### Sparsifying modulation targets

Next we asked whether or not all neuronal parameters in the primary network need to be under modulatory control, or whether a sparse subset would be sufficient. Accuracy remained above the unmodulated baseline across a broad range of sparsity levels, improving as more hidden parameters were made available for modulation. The strongest effect came from retaining all output-layer modulation: with output modulation enabled, performance stayed high even when only a small number of hidden-layer parameters were directly modulated (fig. 5D).

##### Compressing hidden-spike summaries

We also tested compressing the hidden-spike summaries provided to the modulator, either by retaining only the spikes from directly modulated neurons, by grouping hidden neurons that share a modulatory output, or by combining both strategies. These changes had little effect on accuracy (data not shown).

##### Hidden-layer scaling

Finally, we compared modulated and unmodulated models across hidden-layer sizes (fig. 5E). Neuromodulation yields higher accuracy across all tested sizes, with the largest gains for smaller hidden layers.

##### Comparison

We compared all trained configurations by total parameter count. Individual modulated runs vary widely, especially when aggressive sparsity or compression is applied, but the best modulated configurations form a higher-performing frontier than unmodulated SNNs at comparable parameter counts (fig. 5F). This indicates that the performance gains are not simply a consequence of larger models

#### 1.4.2 Regularisation and spike reduction

We reduce spike counts (and therefore total energy consumption in both biological and neuromorphic contexts) by adding a spike penalty to the training objective, varying the strength of this regularisation using the multiplier. Neuromodulation substantially changes the resulting accuracy–spike trade-off compared to an unmodulated system (fig. 6A). Unmodulated SNNs remain substantially lower in accuracy across the full range of spike counts, and fall drastically in performance at low spike counts. By contrast, modulated networks maintain high accuracy across a wide range of spike budgets and remain competitive even when average spike counts are very low.

**Figure 6:**
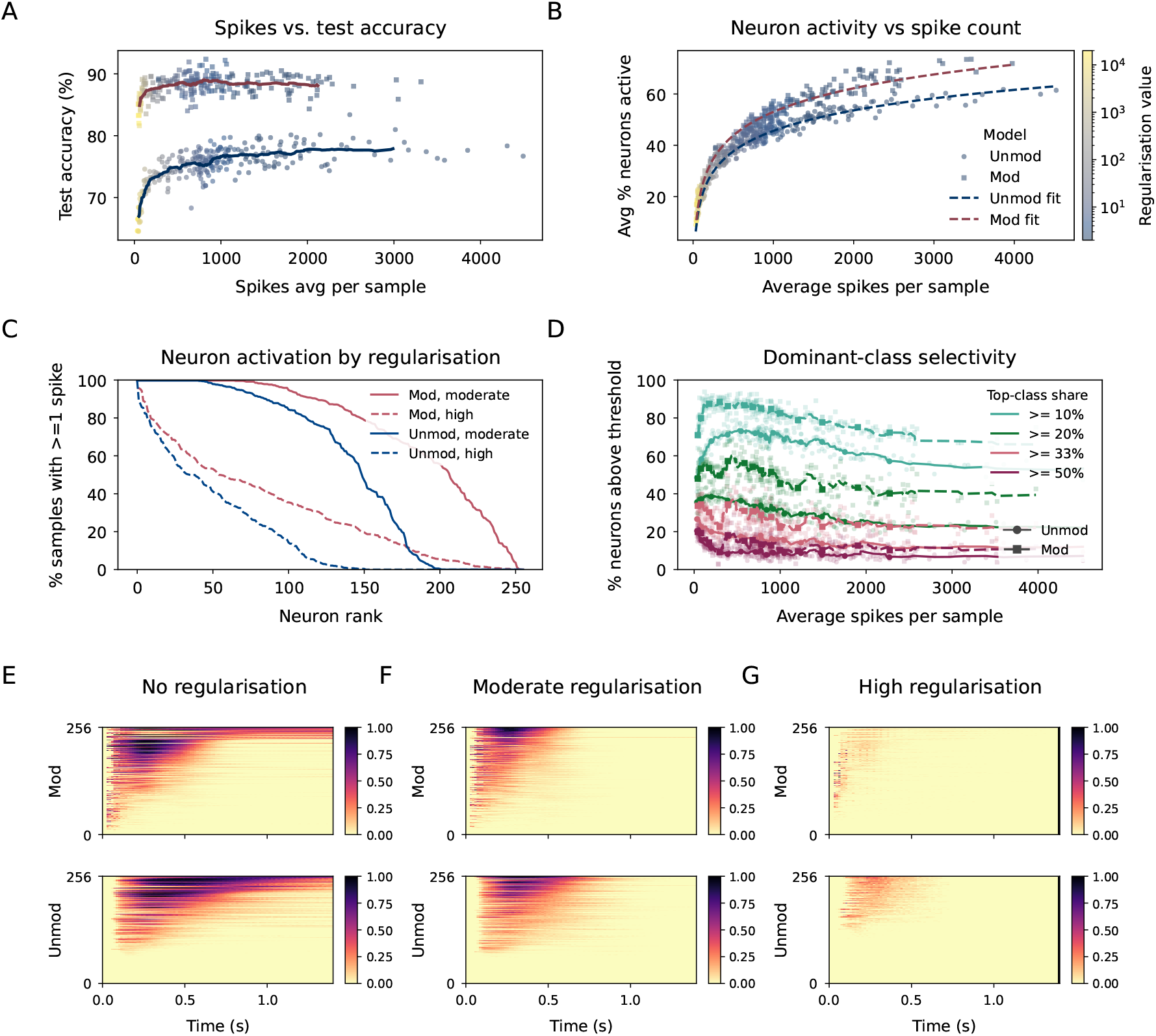
Spike regularisation reveals different activity–accuracy trade-offs in modulated and unmodulated spiking networks. **(A)** Test accuracy as a function of the average number of spikes per sample across a sweep of spike-regularisation strengths. Each point corresponds to one trained model, with colour indicating the regularisation strength. Solid curves show moving averages for the unmodulated SNN and modulated network. **(B)** Hidden-population recruitment under the same regularisation sweep. The y-axis shows the percentage of hidden neurons that fire at least once per sample, plotted against the average spike count per sample. Dashed curves show fitted trends for each model type. For a given spike budget, the modulated network recruits a larger fraction of the hidden population than the unmodulated SNN. **(C)** Per-neuron activation frequency for representative moderate and high regularisation settings. Curves show the percentage of test samples in which each hidden neuron fired at least once, with neurons ranked by activation frequency. Solid lines show moderate regularisation and dashed lines show high regularisation. Modulation maintains activity across more hidden neurons, while the unmodulated SNN develops a larger silent or near-silent population, especially under high regularisation. **(D)** Dominant-class selectivity as a function of spike count. For each trained model, neurons were assigned a dominant output class based on the class that contributed the largest share of their activity. Curves show the percentage of hidden neurons whose dominant class accounts for at least 10%, 20%, 33%, or 50% of their activity. Modulated networks retain a larger population of class-selective neurons across the spike-regularisation sweep. **(E–G)** Example spike-activity maps for no-, moderate-, and high-regularisation. Rows compare modulated and unmodulated networks, with colour indicating normalised spike activity over time and hidden neurons, and are sorted by their centroid.

The modulated networks also use more of the hidden population for a given spike budget (fig. 6B,C), suggesting that modulation promotes distributed sparse codes rather than concentrating most spikes into a small subset of neurons.

At the level of individual neurons, modulation reduces the fraction of completely silent (“dead”) units. Under both moderate and strong regularisation, modulated networks keep a larger fraction of hidden neurons active on at least some test samples, while unmodulated networks develop a larger silent or near-silent tail (fig. 6C). In other words, the effect of spike penalties for unmodulated networks appears to be equivalent to simply switching off some neurons entirely, while with neuromodulation neurons become more context-sensitive.

Spike regularisation also increased dominant-class selectivity in both model types (fig. 6D), consistent with the idea that tighter spike budgets encourage each spike to carry more task-relevant information.

As regularisation increases, both models reduce activity, but they appear to do so in different ways (fig. 6E–G). In the modulated network, spikes become more temporally concentrated earlier in the stimulus, and remain distributed across a relatively broad set of neurons.

## 2 Discussion

We added a simple and flexible learnable model of neuromodulation to a standard spiking neural network. The model essentially allows the network to dynamically adapt the neuronal parameters, and therefore the dynamics and computation of the network, giving it the capability for context-sensitivity. We found that this extra capability gave rise to a substantial improvement in performance across a wide range of tasks and conditions. As an example, it was able to adapt processing in the presence of a time-varying background noise in a manner similar to the “listening in the dips” strategy hypothesised to be a key part of how human listeners are able to hear so well in noisy environments.

Neuromodulation allowed for a considerable reduction in both the space (area) and the energy consumption of the network in both a biological or neuromorphic setting. It did this firstly by reducing the network size needed to attain a certain level of performance. In addition, it allowed the network to achieve high levels of performance with much lower firing rates than an unmodulated network, reducing energy consumption and bandwidth. In general, we found that our modulated networks wasted fewer resources, using less overall neurons and spikes, and using them more efficiently (no dead neurons). These results are highly relevant in both neuroscience and neuromorphic computing, where similar constraints on space and energy are in effect.

For neuromorphic computing, our model has some key properties that will be useful in scaling these models to real-world engineering problems. Firstly, the method is simple to implement. Secondly, it leads to a large gain in performance at a small cost in terms of the number of parameters (and therefore memory). Thirdly, we have shown that it can be used flexibly, either via an artificial or spiking neural network, and at a range of spatial and temporal scales. Finally, it allows us to regularise networks to produce very sparse firing (and therefore energy consumption) without compromising performance. Together, these properties allow us to customise the model for optimal efficiency on each specific device.

In terms of neuroscience, the most direct point of comparison is to experimental studies suggesting a computational role for rapid changes in neuromodulation in sensory processing [Hangya et al., 2015, Bang et al., 2020]. Our results show that it is often more computationally and energetically efficient to add neuromodulation than to add more unmodulated neurons, a strong basis for hypothesising that they play this computational role. Further, these advantages seem to hold across a wide range of tasks and conditions, suggesting they are precisely the sort of robust general purpose mechanisms required in a biological system.

In a more challenging speech in noise task, it discovered an adaptive gain control strategy of ‘listening in the dips’ that has been suggested to be a key part of the human ability to perceive speech in noisy environments [Peters et al., 1998, Lorenzi et al., 2006]. This is an ability that even state-of-the-art machine listening systems still struggle with [O’Shaughnessy, 2024]. Adaptive gain control is a form of rapid attentional mechanism that focuses processing on the most informative parts of a signal. Attention is a key mechanism in the Transformer architecture that heralded the large language model revolution in machine learning [Vaswani et al., 2017], as well as being another hypothesised role of neuromodulation in biology [Avery and Krichmar, 2017, Grossman and Cohen, 2022].

We have shown that neuromodulation can be advantageous for sensory processing in noise via an attentional mechanism. However, this does not exhaust its possible computational roles. Firstly, there may be other sensory processing tasks in which neuromodulation would be advantageous. Secondly, our model uses a simplified general neuromodulatory mechanism in which the output of some neurons can modify various neuronal parameters such as excitability and time constants. This is consistent with known properties of neuromodulators, but not exhaustive [Nadim and Bucher, 2014]. In particular, we do not consider target-cell-type specific effects which are known to be important in the nervous system [Grossman and Cohen, 2022]. In future work, it would be interesting to add additional biophysical constraints and features to our model, and to test these in an even wider range of sensory processing tasks.

Finally, we have not considered in detail how these mechanisms could be implemented across a range of neuromorphic devices. For programmable digital systems such as SpiNNaker [Furber et al., 2014], this would be very straightforward. For analogue, ASIC or hybrid systems, it may be more difficult to implement the full generality of our model of neuromodulation. However, we have found through the Shapley analysis that we do not need all the available mechanisms. Consequently, it is likely that even partial implementations, including sparse modulation that could be integrated via a separate device, would lead to substantial and practical advantages across a wide range of neuromorphic devices.

## 3 Methods

The general structure of the model is as follows (fig. 1A). A neural network is trained to classify a set of inputs represented as spike trains. The network takes as inputs a sequence of spike trains, which are fed into a primary spiking neural network (SNN; sections 3.1.1 and 3.2.2). The output of this network is fed into a linear output layer that is used to classify the inputs with some loss function. The errors from this loss function are backpropagated to update the parameters of the network (section 3.3). In addition, a secondary network (the modulator network) is trained to modulate the parameters of the primary SNN based on the network activity so far. This secondary network can be either an artificial neural network (ANN; sections 3.1.2 and 3.2.3) or another SNN (section 3.2.4). The input to the modulator network is sometimes spatially or temporally compressed (section 3.2.1). The output of the modulator network can be thought of as the release of neuromodulator that goes through three processes: sparse spatial dispersion, interactions between modulator types, and temporal diffusion (section 3.1.3).

All experiments reported in the main figures used a simulation timestep of Δ*t* = 14 ms. Modulator update intervals are therefore reported in milliseconds as *K*Δ*t*, where *K* is the number of simulation steps between neuromodulatory updates. Representative finer-timestep checks gave the same qualitative effect of neuromodulation.

### 3.1 Models

#### 3.1.1 Spiking neurons

The spiking neurons are modeled as leaky integrate-and-fire (LIF) units. Each unit has a membrane potential that evolves over time according to the following differential equations:

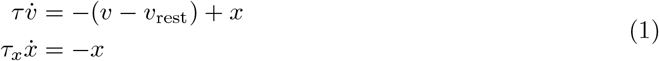

Here, *v* is the membrane potential, *v*_rest_ is the resting potential, *x* is a synaptic variable, *τ* is the membrane time constant, and *τ*_*x*_ is the synaptic time constant. When a spike arrives at the neuron from a synapse with a weight *w*, the following event update is applied:

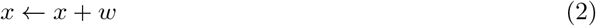

When the membrane potential exceeds a threshold, the neuron emits a spike and resets its potential:

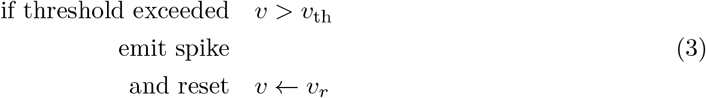

Here *v*_th_ is the threshold, and *v*_*r*_ is the reset value. Note that for simplicity of the implementation, in the code the time constants *τ* and *τ*_*x*_ are not directly used, but instead the parameters *α* and *β* are used, which are defined as:

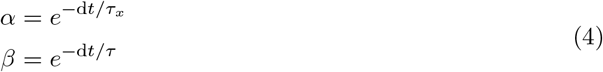

The parameters which can be affected by neuromodulation are *τ*, *τ*_*x*_, *v*_*r*_, *v*_th_ and *v*_rest_.

The output layer of the SNN is non-spiking, and behaves exactly the same as the SNN except for the lack of threshold and reset behaviours, and we additionally fix *v*_rest_ = 0. In this case the parameters that can be affected by neuromodulation are *τ* and *τ*_*x*_.

#### 3.1.2 Artificial neurons

The artificial neurons are modeled as multi-layer perceptrons (MLPs). Each layer of neurons takes as input a vector of activations **x** and computes an output vector **y** using the following equation:

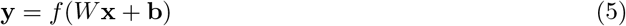

Here, *W* is the weight matrix, **b** is a bias vector, and *f* is an activation function. We use three different activation functions. The rectified linear unit (ReLU):

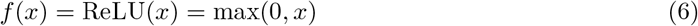

The sigmoid function:

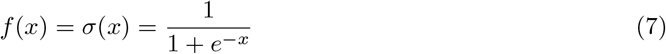

And the hyperbolic tangent function:

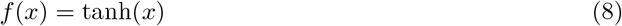

#### 3.1.3 Neuromodulation

Neuromodulation modifies the parameters of a target population of neurons via a process composed of three parts summarised in the equation below, namely spatial dispersion from a release site to the target neuron, interactions between the different types of neuromodulators, and temporal diffusion:

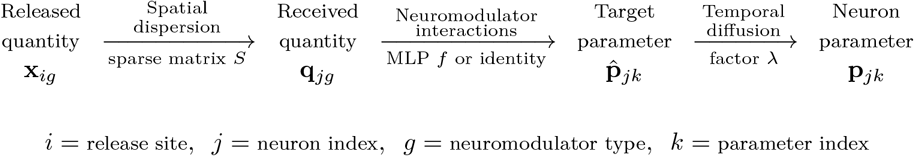

In more detail, the output of the modulator network is a time-varying series of modulator release quantities *x*_*ig*_(*t*) at site *I* ∈{1, …, *M*} (where *M* is the number of release sites) and of type *g* (an integer). When the modulator network is an ANN, *x*_*ig*_(*t*) ∈ℝ, and when it is an SNN, *x*_*ig*_(*t*) ∈ {0, 1} (usually 0).

Each target neuron *j* ∈{1, …, *N*} receives a quantity *q*_*jg*_(*t*) of neuromodulator of type *g*. This quantity is a linear sum over all release sites,

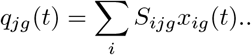

The matrix *S* is sparse and represents the spatial spread from neuromodulator release site *i* to neuron *j*. Throughout the paper we consider different structures for this matrix including 1-to-1 (identity matrix), rectangular windows (overlapping or not), Gaussian windows, or arbitrary sparse matrices.

Each target neuron has *K* parameters *p*_*jk*_(*t*) (where *k* = 1, …, *K*). We do not modify these parameters directly, instead we set a target parameter 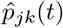 with a slow drift (see next paragraph). The target parameter is a shared function of the vector **q**_*j*_(*t*) of neuromodulator quantities *q*_*jq*_(*t*) at that neuron, and giving as output the new target parameters at that neuron, using one of two formulations:

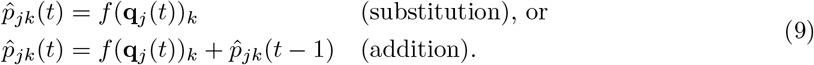

Here *f* is either an identity map when there is one neuromodulator type for each parameter, or an MLP whose parameters are learned (but are shared between all the neurons in the network). This allows for (nonlinear) interactions between different types of neuromodulator. Note that in all cases we clip the output values in the valid range (for example, because time constants cannot be zero or negative).

Note that when the neuromodulator network is an SNN, we always use the additive method, and since we cannot have negative values of *x*_*ig*_(*t*), we include two neuromodulator types for each parameter, one that increases and one that decreases (with learnable size of increase/decrease).

Finally, we model a slow diffusive process where the true parameter value *p*_*jk*_(*t*) slowly drifts to the target value 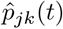 with a diffusion factor *λ* via

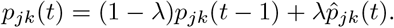

Usually we set *λ* = 1 and this is instantaneous.

As an example, at the start of the paper we consider the situation where we have one neuromodulator release site for each neuron, and one neuromodulator type for each parameter, making both the dispersion map and the interactions map into identity functions, and the diffusion parameter *λ* = 1 for an instantaneous effect. Next, when we talk about parameter grouping with grouping factor *G* that corresponds to setting *M* = *N/G* and setting

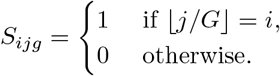

We implement the overlapping fan-out similarly but with an overlap. For Gaussian spatial fan-out, release sites were placed every *G* hidden neurons, with site *i* centred at *i*^*′*^ = *iG* + (*G*−1)*/*2. The effect of release site *i* on hidden neuron *j* was proportional to

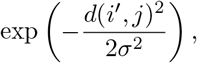

where *d*(·, ·) is circular distance along the hidden-neuron index and *σ* is the spread parameter. Weights below 0.05 were set to zero. The remaining weights determined how strongly each release site modulated each hidden neuron, so nearby neurons received stronger modulation and distant neurons received weak or no modulation.

### 3.2 Architectures

The overall architecture is shown in fig. 1A. The input spikes are defined by the dataset. The primary SNN receives inputs as spikes from these dataset-defined input spikes, and also from itself (the primary SNN is recurrent). It has its parameters modulated by the modulator network, and it outputs a time-varying set of membrane potentials of its output layer. The modulator network receives activity from the input spikes, primary SNN and output membrane potentials (possibly compressed, see below). Internally, the modulator network is either an ANN or an SNN.

The ANN modulator network runs every *K* time steps (where *K* is the modulation interval, a hyperparameter we vary). The SNN modulator network runs every time step, but certain inputs to this network are only updated every *K* time steps, and the outputs are accumulated over *K* time steps and applied at the end of each interval.

We now describe each of these pieces in more detail.

#### 3.2.1 Inputs to modulator network

For both the ANN and SNN modulator, we provide activities and current parameter values as inputs to the network as a concatenated vector. In some cases, we also compress these inputs to reduce dimensionality. We describe the inputs and compression scheme below. First, we start with some notation to simplify this description:

- *X*_*it*_ ∈ {0, 1} are the input spikes for channel *i* at time *t*
- *H*_*it*_ ∈ {0, 1} are the hidden layer spikes
- *V*_*it*_ ∈ ℝ is the output membrane potential
- *P*_*ikt*_ ∈ ℝ is parameter *k* of hidden layer

In the uncompressed scheme with a time interval *K* = 1, the input to the modulators is just a vector concatenating all the values for a fixed *t* (flattening indices *i* and *k* of *P*). However, to reduce the size of the input, we compress in various ways. First, we can spatially compress. Let *Y*_*it*_ be an array of inputs with a spatial dimension *i* and a temporal dimension *t*. We compute the compressed version 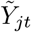by summing over blocks of indices *i* of width *w*:

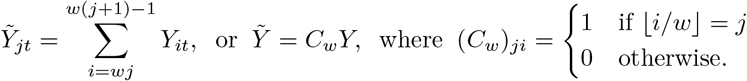

We can do temporal compression by just doing the same thing over the second, temporal dimension instead of the first, this time with temporal width *K* to get 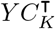 . We can do both spatial and temporal compression then simply by 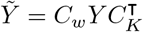.

In addition, we consider filtering: for the hidden layer, *F* [*H*] has all the rows *i* that correspond to neurons that are not modulated removed. Finally, we consider deduplication: *D*[*P*] has all copies of the parameters that have identical values (e.g. because of grouping) reduced to a single value. With this in mind, the compressed inputs to the ANN modulator with spatial grouping *w*, temporal grouping *K* are:

- 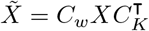input: s undergo spatial and temporal compression
- 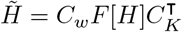: hidden units undergo filtering and then spatial and temporal compression
- 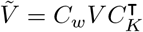: output potentials undergo spatial and temporal compression
- 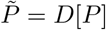: parameter values undergo deduplication

The concatenated vector of all these is provided at every time step of the ANN modulator, which is every *K* time steps of the primary SNN. For the SNN modulator, it has to run at every time step so we do not do temporal compression. In this case we get:

- 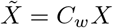
- 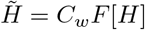
- 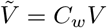
- 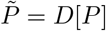

*X, H* and *V* are provided at every SNN time step, and *P* is provided as a set of input currents that are held fixed over the modulation interval.

In the Shapley analysis for both ANN and SNN modulators, some of these component vectors will not be included in the input vector to the modulator.

#### 3.2.2 Primary SNN

The primary SNN consists of a single recurrent hidden layer of leaky integrate-and-fire neurons, connected to a non-recurrent readout layer of leaky but non-spiking neurons (identical equations but no threshold and reset mechanism). Every connection (input to hidden, hidden to hidden and hidden to output) is fully connected and represented by a corresponding weight matrix.

#### 3.2.3 Modulator ANN

The ANN modulator network consists of a two-layer multi-layer perceptron (MLP). The first layer is a linear layer with a ReLU activation function, and the second layer is linear with a sigmoidal activation function (in the case of the substitution method, to keep parameter values within the desired range) or tanh activation function (in the case of the addition method, to allow values to be positive or negative but bounded). For the substitution method an offset is applied to the output for some parameters (+0.5 for threshold, -0.5 for rest and reset).

#### 3.2.4 Modulator SNN

The SNN modulator network consists of a single recurrent layer of leaky integrate-and-fire neurons. The network receives the same inputs as for the ANN modulator, but in this case they are treated as input currents to the neurons (for spiking inputs this is identical to treating them as input spikes). Each output neuron corresponds to a positive or negative change to a specific parameter value. All weight matrices and the size of the output changes are learnable.

#### 3.2.5 Interaction MLP

For the neuromodulator release model, the modulator outputs a vector **q**_*j*_ of neuromodulator quantities for each target neuron or neuron group, rather than one value for each modulated parameter. These quantities are converted into parameter effects by a shared interaction MLP:

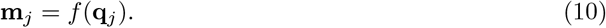

A separate MLP was used for each population. The hidden-layer MLP produced five outputs corresponding to *α, β, v*_th_, *v*_*r*_ and *v*_rest_, while the readout-layer MLP produced two outputs corresponding to *α* and *β*. The same MLP parameters were shared across all target neurons within a population, allowing the neuromodulator types to have consistent interactions throughout that population. Each interaction MLP consisted of two hidden layers of 64 units followed by a linear output layer. For the reported neuromodulator-channel experiments, the interaction map used linear hidden transformations, with no nonlinear activation between hidden layers. The output layer used a sigmoid activation for substitution-based modulation and a tanh activation for addition-based modulation. The resulting outputs were mapped to the corresponding parameter ranges and clipped using the same bounds as in the direct modulation model. All weights and biases of the interaction MLPs were learned jointly with the modulator network.

### 3.3 Training

All networks were trained using surrogate gradient descent [Neftci et al., 2019], which, in the backwards pass only, replaces the non-differentiable spike function with a smoothed sigmoid to enable backpropagation. Specifically, in the forward pass we use the Heaviside function

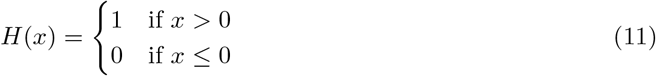

in the threshold equation *S* = *H*(*v* − *v*_th_), where *S* = 1 indicates a spike. When computing gradients in the backwards pass, we replace *H*^*′*^(*x*) with 1*/*(|*x*| + 1)2 as in Zenke and Ganguli [2018].

We used the Adam optimiser [Kingma and Ba, 2017] with dataset-specific initial learning rates listed in table 1. The following parameters are learnable. For the primary SNN: the input-to-hidden weight matrix, the hidden-to-output weight matrix, the hidden-to-hidden recurrent weight matrix, and the initial value of each parameter for each neuron (membrane and synaptic time constants, threshold, reset, rest). For the ANN modulator: the weights and biases of the modulator network. For the SNN modulator: the modulator layer weights, any recurrent weights, the modulator neuron parameters, and the learned output-effect scales. In interacting neuromodulator type experiments, the modulator outputs neuromodulator quantities rather than direct parameter effects; the weights and biases of the shared interaction MLPs that map these quantities to hidden and readout-layer parameter effects are also learned jointly.

**Table 1.** Model architecture and training parameters. The top rows give dataset-specific input, hidden, output and learning-rate settings; the lower rows give shared neuron constants, clipping ranges and training settings.

| Parameter | SHD | SSC | DVS | Description |
| --- | --- | --- | --- | --- |
| Input neurons | 700 | 700 | 4096 | No. of preprocessed input units |
| Hidden spiking neurons | 256 | 256 | 256 | Size of hidden LIF layer |
| Output neurons | 20 | 35 | 11 | Number of classes |
| Learning rate | $2 \times 10^{-4}$ | $2 \times 10^{-5}$ | $2 \times 10^{-4}$ | Initial optimizer learning rate |
| $\tau$ | | 20 ms | | Membrane time constant |
| $\tau_x$ | | 10 ms | | Synaptic time constant |
| $v_{th}$ | | 1 V | | Spiking threshold |
| $v_{rest}$ | | 0 mV | | Resting potential |
| $v_r$ | | 0 mV | | Reset potential |
| $\rho$ | | 100 | | Surrogate-gradient steepness |
| Time constant min | | 1 ms | | Minimum $\tau, \tau_x$ |
| Time constant max | | 200 ms | | Maximum $\tau, \tau_x$ |
| Threshold min | | 0.5 V | | Minimum $v_{th}$ |
| Threshold max | | 1.5 V | | Maximum $v_{th}$ |
| Reset & rest min | | -0.5 V | | Minimum $v_{rest}, v_r$ |
| Reset & rest max | | 0.5 V | | Maximum $v_{rest}, v_r$ |
| Optimizer |  | Adam |  | Optimization algorithm |
| Batch size |  | 64 |  | Training batch size |
| Early stopping patience |  | 10 epochs |  | Validation-loss patience used for checkpoint selection |

For each dataset, a validation set was held out from the original training set. Training proceeded in two phases:

1. The primary SNN was first trained without modulation. Training stopped if the validation loss did not improve for 10 consecutive epochs, and the checkpoint with the lowest validation loss was retained. This checkpoint was used both as the unmodulated baseline and to initialise the modulated network.
2. The primary SNN and modulator network were then trained jointly using the same early-stopping and checkpoint-selection procedure.

Hyperparameters were selected using the training and validation sets. The test set was reserved for final evaluation of the selected checkpoint.

The standard training loss consisted of three terms: a task loss and two spike-rate regularisation terms.

Let *v*_*i*_(*t*) be the membrane potential of output neuron *i* at time *t*. We define the output vector

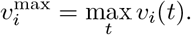

The output logits are then set to

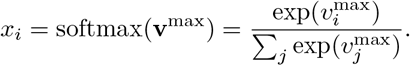

The task loss is then the cross-entropy loss on these logit values.

The regularisation terms keep spike rates within reasonable ranges and help to avoid instability in training. There are two terms. The first term is proportional to the sum over neuron and batch indices of 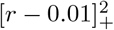, where *r* is the neuron’s firing rate and [*x*]_+_ = max(*x*, 0). The second term is proportional to the sum over the batch of 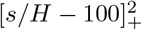, where *s* is the total hidden spike count for a sample and *H* is the number of hidden neurons. For the spike-regularisation sweeps in fig. 6, we added explicit spike penalties and swept their multiplier. We report average hidden spikes per sample, the percentage of hidden neurons active at least once per sample, per-neuron activation frequency across test samples, and dominant-class selectivity computed from each neuron’s class-conditioned activity.

Parameter values were clipped to stay within biologically plausible ranges: time constants were clipped between 1-200 ms; thresholds were clipped between 0.5 and 1.5.

### 3.4 Analyses

For all plots, unless otherwise stated, points show individual trained models or means across independent runs, and shaded regions show one standard deviation across runs. Difference panels report modulated minus unmodulated accuracy, except where explicitly stated otherwise.

For the time-resolved natural-noise analysis in fig. 2C–E, the background-noise envelope was computed as root-mean square (RMS) amplitude in 14 ms frames, matching the simulation timestep. For fig. 2C, the background-noise envelope and average spike activity were smoothed with a 555-ms moving-average window before normalisation and plotting. Spike activity was averaged across evaluated trials. Lagged noise–spike coupling was measured using Pearson correlation between the noise envelope and spike activity after shifting one trace relative to the other; the plotted curve reports correlation as a function of lag, with the strongest negative correlation at -14 ms. For the scatter plot, time windows were sampled from the aligned traces and both noise and spike activity were normalised before fitting the linear trend.

For the sinusoidal-noise hidden/input analysis in fig. 3H, input and hidden spike probabilities were averaged across test samples and neurons at each time bin. We then divided mean hidden activity by mean input activity to obtain a hidden/input activity ratio. This ratio was correlated with the known sinusoidal noise envelope over the 0–800 ms analysis window. The lower panel reports the difference between unmodulated and modulated correlations, computed as unmodulated minus modulated.

For the Shapley analyses in fig. 5B,C, modulator inputs and outputs were grouped into blocks corresponding to input spikes, hidden spikes, output potentials, and parameter groups. Block importance was estimated using a Monte Carlo permutation estimator of the Shapley value [Castro et al., 2009, Mitchell et al., 2022]. For each random ordering, blocks were enabled one at a time from an all-disabled baseline, and the marginal change in accuracy was accumulated. Blocks were then ranked by absolute Shapley contribution. The cumulative plots add blocks in this rank order and report test accuracy across five runs.

Parameter counts in fig. 5F were computed as total trainable model parameters, including both the primary SNN and the modulator network. The plotted frontier is the 95th-percentile accuracy envelope over sliding windows in log-parameter space. It therefore summarises the best-performing configurations at each model scale, rather than the mean over all configurations.

For the spike-regularisation analysis in fig. 6, regularisation strength refers to the multiplier applied to the spike-rate penalty during training, swept jointly for the unmodulated and modulated models. Figure 6A plots checkpoint test accuracy against average hidden spikes per sample. Figure 6B measures hidden-population recruitment by replaying the test set and computing the percentage of hidden neurons that fired at least once per sample. Figure 6C plots, for each hidden neuron, the percentage of test samples in which that neuron fired at least once, with neurons ranked by activation frequency. Figure 6D measures dominant-class selectivity: for each neuron, class-conditioned activity was accumulated across the test set, the class contributing the largest share of that neuron’s activity was identified, and the plot reports the percentage of neurons whose dominant-class share exceeded each threshold.

### 3.5 Datasets

#### 3.5.1 Standard datasets

We evaluated our models on three standard datasets:

- **Spiking Heidelberg Digits (SHD):** A clean audio digit dataset of ten spoken digits by 12 distinct speakers in English and German, processed via a model of the cochlea into 700-channel spike trains [Cramer et al., 2022]. The dataset consists of 20 classes, 8156 training samples, and 2264 testing samples. Two of the speakers are only present in the test set.
- **Spiking Speech Commands (SSC):** A more complex and noisy speech dataset with 35 classes, many different speakers, and diverse recording conditions from the Google Speech Commands dataset [Warden, 2018] processed into spike trains in the same way as SHD. The dataset consists of 35 classes, 75466 training samples, 9981 validation samples, and 20382 testing samples.
- **DVS128 Gestures:** An event-based video gesture dataset captured using a 128 × 128 pixel dynamic vision sensor [Amir et al., 2017]. Contains 11 gestures from 29 subjects under 3 illumination conditions. 23 subjects are used for the training set, and 6 for the testing set. In total, there are 1342 samples in the dataset. The model input used in our experiments is the preprocessed event representation listed in table 1.

#### 3.5.2 Speech-in-noise dataset

For the speech in noise experiments we used the following process [AlKilany and Goodman, 2026]. We took the original target audio files used in the SHD dataset, added either sinusoidally amplitude-modulated (SAM) noise or natural noise, and then converted into spike trains using the same technique as in Cramer et al. [2022].

Note that after correspondence with the authors, we found that the code available online at https://github.com/electronicvisions/lauscher does not exactly correspond to what was used in the paper. To reproduce their results we had to modify their code to reduce the size of the Hanning window to a 30 ms ramp-up and ramp-down time.

The SAM noise is simply computed by multiplying a white noise (Gaussian distributed samples) with an envelope

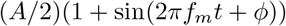

where *A* is the amplitude, *f*_*m*_ is the modulation frequency and *ϕ* is the starting phase. For the plots in the paper, we set *ϕ* = 0 to make it possible to visualise the effect, but in the training and testing we randomise *ϕ* uniformly in [0, 2*π*).

For the natural-noise experiments, we used cafe background recordings from the noisy speech dataset of Reddy et al. [2019]. We used randomly selected segments of the background noise with duration equal to the target sound duration.

In the case of both types of noise, we use a range of signal-to-noise ratios (SNRs). We use the target sounds calibrated in the same way as in Cramer et al. [2022], only modifying the amplitude of the noise to vary the SNR.

### 3.6 Reproducibility and code

All code and experiments were implemented in PyTorch [Paszke et al., 2019]. The code and reproducibility instructions are available at https://github.com/abdalalkilani/NeuromodulationSNN.

The Noisy SHD dataset is available from Zenodo [AlKilany and Goodman, 2026] with instructions on use available at https://github.com/abdalalkilani/Noisy-SHD.

## A Supplementary materials

### A.1 Architecture parameters

## References

S. B. Laughlin, R. R. de Ruyter van Steveninck, and J. C. Anderson. The metabolic cost of neural information. Nature Neuroscience, 1(1):36–41, May 1998. ISSN 1097-6256. doi:10.1038/236.

David Attwell and Simon B Laughlin. An energy budget for signaling in the grey matter of the brain. Journal of Cerebral Blood Flow & Metabolism, 21(10):1133–1145, 2001.

Peter Lennie. The cost of cortical computation. Current biology, 13(6):493–497, 2003.

Catherine D. Schuman, Shruti R. Kulkarni, Maryam Parsa, J. Parker Mitchell, Prasanna Date, and Bill Kay. Opportunities for neuromorphic computing algorithms and applications. Nature Computational Science, 2(1):10–19, January 2022. ISSN 2662-8457. doi:10.1038/s43588-021-00184-y. URL https://www.nature.com/articles/s43588-021-00184-y. Publisher: Nature Publishing Group.

Emre O. Neftci, Hesham Mostafa, and Friedemann Zenke. Surrogate Gradient Learning in Spiking Neural Networks: Bringing the Power of Gradient-Based Optimization to Spiking Neural Networks. IEEE Signal Processing Magazine, 36(6):51–63, November 2019. ISSN 1558-0792. doi:10.1109/MSP.2019.2931595. Conference Name: IEEE Signal Processing Magazine.

Guillaume Bellec, Franz Scherr, Anand Subramoney, Elias Hajek, Darjan Salaj, Robert Legenstein, and Wolfgang Maass. A solution to the learning dilemma for recurrent networks of spiking neurons. Nature communications, 11(1):3625, 2020.

Jason K. Eshraghian, Max Ward, Emre O. Neftci, Xinxin Wang, Gregor Lenz, Girish Dwivedi, Mohammed Bennamoun, Doo Seok Jeong, and Wei D. Lu. Training Spiking Neural Networks Using Lessons From Deep Learning. Proceedings of the IEEE, 111(9):1016–1054, September 2023. ISSN 1558-2256. doi:10.1109/JPROC.2023.3308088. URL https://ieeexplore.ieee.org/document/10242251.

Dhireesha Kudithipudi, Catherine Schuman, Craig M. Vineyard, Tej Pandit, Cory Merkel, Rajkumar Kubendran, James B. Aimone, Garrick Orchard, Christian Mayr, Ryad Benosman, Joe Hays, Cliff Young, Chiara Bartolozzi, Amitava Majumdar, Suma George Cardwell, Melika Payvand, Sonia Buckley, Shruti Kulkarni, Hector A. Gonzalez, Gert Cauwenberghs, Chetan Singh Thakur, Anand Subramoney, and Steve Furber. Neuromorphic computing at scale. Nature, 637(8047): 801–812, January 2025. ISSN 1476-4687. doi:10.1038/s41586-024-08253-8. URL https://www.nature.com/articles/s41586-024-08253-8. Publisher: Nature Publishing Group.

Mike Davies, Andreas Wild, Garrick Orchard, Yulia Sandamirskaya, Gabriel A. Fonseca Guerra, Prasad Joshi, Philipp Plank, and Sumedh R. Risbud. Advancing Neuromorphic Computing With Loihi: A Survey of Results and Outlook. Proceedings of the IEEE, 109(5):911–934, May 2021. ISSN 1558-2256. doi:10.1109/JPROC.2021.3067593. URL https://ieeexplore.ieee.org/document/9395703.

Nicolas Perez-Nieves, Vincent C. H. Leung, Pier Luigi Dragotti, and Dan F. M. Goodman. Neural heterogeneity promotes robust learning. Nature Communications, 12(1):5791, October 2021. ISSN 2041-1723. doi:10.1038/s41467-021-26022-3. URL . https://www.nature.com/articles/s41467-021-26022-3. Number: 1 Publisher: Nature Publishing Group.

David Dahmen, Axel Hutt, Giacomo Indiveri, Ann Kennedy, Jeremie Lefebvre, Luca Mazzucato, Adilson E. Motter, Rishikesh Narayanan, Melika Payvand, Henrike Planert, and Richard Gast. How heterogeneity shapes dynamics and computation in the brain. Neuron, 114(5):804–819, March 2026. ISSN 0896-6273. doi:10.1016/j.neuron.2025.11.023. URL https://www.sciencedirect.com/science/article/pii/S0896627325008967.

Guillaume Bellec, Darjan Salaj, Anand Subramoney, Robert Legenstein, and Wolfgang Maass. Long short-term memory and Learning-to-learn in networks of spiking neurons. In Advances in Neural Information Processing Systems, volume 31. Curran Associates, Inc., 2018. URL https://papers.nips.cc/paper_files/paper/2018/hash/c203d8a151612acf12457e4d67635a95-Abstract.html.

Darjan Salaj, Anand Subramoney, Ceca Kraisnikovic, Guillaume Bellec, Robert Legenstein, and Wolfgang Maass. Spike frequency adaptation supports network computations on temporally dispersed information. eLife, 10:e65459, July 2021. ISSN 2050-084X. doi:10.7554/eLife.65459. URL https://doi.org/10.7554/eLife.65459.

Pengfei Sun, Yansong Chua, Paul Devos, and Dick Botteldooren. Learnable axonal delay in spiking neural networks improves spoken word recognition. Frontiers in Neuroscience, 17:1275944, 2023.

Pengfei Sun, Jascha Achterberg, Zhe Su, Dan F. M. Goodman, and Danyal Akarca. Exploiting heterogeneous delays for efficient computation in low-bit neural networks, October 2025. URL http://arxiv.org/abs/2510.27434. arXiv:2510.27434 [cs.NE].

Pengfei Sun, Zhe Su, Jascha Achterberg, Giacomo Indiveri, Dan F. M. Goodman, and Danyal Akarca. Algorithm–hardware co-design of neuromorphic networks with dual memory pathways. Nature Machine Intelligence, 8(6):901–912, June 2026. ISSN 2522-5839. doi:10.1038/s42256-026-01255-3. URL https://www.nature.com/articles/s42256-026-01255-3.

Dzmitry Bahdanau, Kyunghyun Cho, and Yoshua Bengio. Neural machine translation by jointly learning to align and translate. arXiv preprint arXiv:1409.0473, 2014.

Ashish Vaswani, Noam Shazeer, Niki Parmar, Jakob Uszkoreit, Llion Jones, Aidan N. Gomez, Lukasz Kaiser, and Illia Polosukhin. Attention Is All You Need. arXiv:1706.03762 [cs], December 2017. URL http://arxiv.org/abs/1706.03762. arXiv: 1706.03762.

Farzan Nadim and Dirk Bucher. Neuromodulation of Neurons and Synapses. Current opinion in neurobiology, 0:48–56, December 2014. ISSN 0959-4388. doi:10.1016/j.conb.2014.05.003. URL https://www.ncbi.nlm.nih.gov/pmc/articles/PMC4252488/.

Jie Mei, Eilif Muller, and Srikanth Ramaswamy. Informing deep neural networks by multiscale principles of neuromodulatory systems. Trends in Neurosciences, 45(3):237–250, March 2022. ISSN 0166-2236. doi:10.1016/j.tins.2021.12.008. URL https://www.sciencedirect.com/science/article/pii/S0166223621002563.

Kenji Doya. Metalearning and neuromodulation. Neural Networks, 15(4):495–506, June 2002. ISSN 0893-6080. doi:10.1016/S0893-6080(02)00044-8. URL https://www.sciencedirect.com/science/article/pii/S0893608002000448.

Cooper D. Grossman and Jeremiah Y. Cohen. Neuromodulation and Neurophysiology on the Timescale of Learning and Decision-Making. Annual Review of Neuroscience, 45:317–337, July 2022. ISSN 1545-4126. doi:10.1146/annurev-neuro-092021-125059.

Balázs Hangya, Sachin P. Ranade, Maja Lorenc, and Adam Kepecs. Central Cholinergic Neurons Are Rapidly Recruited by Reinforcement Feedback. Cell, 162(5):1155–1168, August 2015. ISSN 0092-8674, 1097-4172. doi:10.1016/j.cell.2015.07.057. URL https://www.cell.com/cell/abstract/S0092-8674(15)00973-3. Publisher: Elsevier.

Dan Bang, Kenneth T. Kishida, Terry Lohrenz, Jason P. White, Adrian W. Laxton, Stephen B. Tatter, Stephen M. Fleming, and P. Read Montague. Sub-second Dopamine and Serotonin Signaling in Human Striatum during Perceptual Decision-Making. Neuron, 108(5):999–1010.e6, December 2020. ISSN 0896-6273. doi:10.1016/j.neuron.2020.09.015. URL https://www.sciencedirect.com/science/article/pii/S0896627320307157.

Jeffrey L. Krichmar. A biologically inspired action selection algorithm based on principles of neuromodulation. In The 2012 International Joint Conference on Neural Networks (IJCNN), pages 1–8, June 2012. doi:10.1109/IJCNN.2012.6252633. URL https://ieeexplore.ieee.org/abstract/document/6252633. ISSN: 2161-4407.

Luka Ribar and Rodolphe Sepulchre. Neuromodulation of neuromorphic circuits. IEEE Transactions on Circuits and Systems I: Regular Papers, 66(8):3028–3040, 2019.

Samuel Schmidgall, Julia Ashkanazy, Wallace Lawson, and Joe Hays. Spikepropamine: Differentiable plasticity in spiking neural networks. Frontiers in neurorobotics, 15:629210, 2021.

Benjamin Cramer, Yannik Stradmann, Johannes Schemmel, and Friedemann Zenke. The Heidelberg Spiking Data Sets for the Systematic Evaluation of Spiking Neural Networks. IEEE Transactions on Neural Networks and Learning Systems, 33(7):2744–2757, July 2022. ISSN 2162-2388. doi:10.1109/TNNLS.2020.3044364. URL https://ieeexplore.ieee.org/document/9311226.

Arnon Amir, Brian Taba, David Berg, Timothy Melano, Jeffrey McKinstry, Carmelo Di Nolfo, Tapan Nayak, Alexander Andreopoulos, Guillaume Garreau, Marcela Mendoza, Jeff Kusnitz, Michael Debole, Steve Esser, Tobi Delbruck, Myron Flickner, and Dharmendra Modha. A Low Power, Fully Event-Based Gesture Recognition System. In 2017 IEEE Conference on Computer Vision and Pattern Recognition (CVPR), pages 7388–7397, July 2017. doi:10.1109/CVPR.2017.781. URL https://ieeexplore.ieee.org/document/8100264. ISSN: 1063-6919.

AbdelQader AlKilany and Dan Goodman. Noisy SHD: Spiking Heidelberg Digits with sinusoidal and natural noise, 2026. URL 10.5281/zenodo.21313625.

Robert W. Peters, Brian C. J. Moore, and Thomas Baer. Speech reception thresholds in noise with and without spectral and temporal dips for hearing-impaired and normally hearing people. The Journal of the Acoustical Society of America, 103(1):577–587, January 1998. ISSN 0001-4966. doi:10.1121/1.421128. URL https://doi.org/10.1121/1.421128.

Christian Lorenzi, Gaëtan Gilbert, Héloïse Carn, Stéphane Garnier, and Brian C. J. Moore. Speech perception problems of the hearing impaired reflect inability to use temporal fine structure. Proceedings of the National Academy of Sciences, 103(49):18866–18869, December 2006. doi:10.1073/pnas.0607364103. URL https://www.pnas.org/doi/abs/10.1073/pnas.0607364103. Publisher: Proceedings of the National Academy of Sciences.

Douglas O’Shaughnessy. Trends and developments in automatic speech recognition research. Computer Speech & Language, 83:101538, January 2024. ISSN 0885-2308. doi:10.1016/j.csl.2023.101538. URL https://www.sciencedirect.com/science/article/pii/S0885230823000578.

Michael C. Avery and Jeffrey L. Krichmar. Neuromodulatory Systems and Their Interactions: A Review of Models, Theories, and Experiments. Frontiers in Neural Circuits, 11, December 2017. ISSN 1662-5110. doi:10.3389/fncir.2017.00108. URL https://www.frontiersin.org/journals/neural-circuits/articles/10.3389/fncir.2017.00108/full. Publisher: Frontiers.

Steve B. Furber, Francesco Galluppi, Steve Temple, and Luis A. Plana. The spinnaker project. Proceedings of the IEEE, 102(5):652–665, 2014. doi:10.1109/JPROC.2014.2304638.

Friedemann Zenke and Surya Ganguli. SuperSpike: Supervised learning in multilayer spiking neural networks. Neural Computation, 2018. ISSN 1530888X. doi:10.1162/neco_a_01086.

Diederik P. Kingma and Jimmy Ba. Adam: A Method for Stochastic Optimization, January 2017. URL http://arxiv.org/abs/1412.6980. arXiv:1412.6980 [cs].

Javier Castro, Daniel Gómez, and Juan Tejada. Polynomial calculation of the shapley value based on sampling. Computers & operations research, 36(5):1726–1730, 2009.

Rory Mitchell, Joshua Cooper, Eibe Frank, and Geoffrey Holmes. Sampling permutations for shapley value estimation. Journal of Machine Learning Research, 23(43):1–46, 2022.

Pete Warden. Speech Commands: A Dataset for Limited-Vocabulary Speech Recognition, April 2018. URL http://arxiv.org/abs/1804.03209. arXiv:1804.03209 [cs].

Chandan KA Reddy, Ebrahim Beyrami, Jamie Pool, Ross Cutler, Sriram Srinivasan, and Johannes Gehrke. A scalable noisy speech dataset and online subjective test framework. Proc. Interspeech 2019, pages 1816–1820, 2019.

Adam Paszke, Sam Gross, Francisco Massa, Adam Lerer, James Bradbury, Gregory Chanan, Trevor Killeen, Zeming Lin, Natalia Gimelshein, Luca Antiga, Alban Desmaison, Andreas Köpf, Edward Yang, Zach DeVito, Martin Raison, Alykhan Tejani, Sasank Chilamkurthy, Benoit Steiner, Lu Fang, Junjie Bai, and Soumith Chintala. PyTorch: An Imperative Style, High-Performance Deep Learning Library, December 2019. URL http://arxiv.org/abs/1912.01703. arXiv:1912.01703 [cs, stat].

